# Lifelong self-renewal and competition shape the virtual memory CD8 T-cell compartment during aging

**DOI:** 10.64898/2026.06.07.730686

**Authors:** Bo Chin Chiu

## Abstract

Primary CD8^+^ T-cell responses decline with age, increasing susceptibility to novel infections. Virtual memory (VM) CD8^+^ T cells are memory-phenotype cells that respond rapidly to infection and undergo marked expansion with age. Although aged VM cells exhibit functional impairment, cellular senescence, and clonal expansion, the mechanisms responsible for these changes remain poorly understood. Here, we show that the VM-cell compartment in aged mice is derived predominantly from cells generated early in life and maintained through continuous self-renewal. Over time, resident VM cells acquire increased competitive fitness, resulting in progressive enrichment of the resident population and exclusion of newly generated VM cells. Consequently, unlike the naïve CD8^+^ T-cell compartment, which is continuously replenished throughout life, the VM-cell compartment becomes dominated by long-lived resident cells. Depletion of resident VM cells resets the aged VM-cell compartment and permits expansion of newly generated VM cells. These findings identify lifelong self-renewal and competition as key mechanisms shaping VM-cell aging.

## Introduction

Steady-state CD8^+^ T cells can be broadly divided into naïve and memory-phenotype (MP) populations [1], with the latter further subdivided into virtual memory (VM) and true memory (TM) subsets [2]. Unlike TM cells, which arise following exposure to foreign antigens, VM cells develop under steady-state conditions from a distinct lineage of autoreactive CD8 single-positive thymocytes [3]. The developmental pathways of VM and naïve CD8^+^ T cells diverge in the thymus, where EOMES functions as a lineage-determining transcription factor for VM-cell development [3]. Following thymic commitment, VM cells undergo post-thymic maturation and acquire their characteristic phenotype, including high expression of the IL-2/IL-15 receptor β chain (CD122) and low expression of CD49d [2-4].

A distinctive feature of VM cells is their ability to respond rapidly to infection [5-8]. VM cells exhibit innate-like effector functions and can contribute to primary immune responses against pathogens. Childhood infections often establish lifelong immunological memory, and pathogen-specific memory T cells generated early in life generally remain functional into old age [9, 10]. In contrast, aged individuals are more susceptible to novel infections, in part because primary CD8^+^ T-cell responses decline with age [9, 11]. Both naïve and VM cells contribute to responses against new infections [5-7], yet these populations appear to age differently. The number of naïve CD8^+^ T cells declines progressively with age, whereas the VM-cell compartment undergoes dramatic expansion. Despite this numerical increase, aged VM cells exhibit functional impairment, features of cellular aging, and clonal expansion [5, 6, 12-16].

Although age-associated changes in VM cells have been well documented, the mechanisms responsible for these changes remain poorly understood. In the present study, we examined the origin, maintenance, and long-term dynamics of the VM-cell compartment during aging. We show that most VM cells present in aged mice originate early in life and are maintained through continuous self-renewal. Over time, the VM-cell compartment becomes enriched for cells with increased competitive fitness and progressively excludes newly generated VM cells. These findings identify lifelong self-renewal and competition among resident VM cells as key mechanisms shaping the aged VM-cell compartment.

## Results

### VM cells in aged mice arise predominantly early in life

A peripheral CD8^+^ T-cell compartment is established shortly after birth. During this period, the frequency of VM cells is high and comparable to that observed in aged mice because CD8^+^ T cells generated during the neonatal period are substantially more likely to adopt the VM fate than those generated in adulthood [17, 18]. This difference is thought to reflect, at least in part, the unique properties of fetal hematopoietic stem cells (HSCs). Whether age-associated changes in HSCs influence the propensity of CD8^+^ T cells to differentiate into VM cells remains unclear.

To directly compare the VM-generating potential of young and aged HSCs, congenically marked bone marrow (BM) cells from young (CD45.1^+^ CD45.2^+^) and aged (CD45.1^−^CD45.2^+^) donors were co-transferred into lethally irradiated young recipients (CD45.1^+^ CD45.2^−^) (Figure 1A). Donor-derived cells were analyzed 15 months after transplantation. Unexpectedly, a substantial population of host-derived cells remained within the peripheral CD8^+^ T-cell compartment (Figure 1B). This persistence was unlikely to reflect incomplete myeloablation because host-derived cells were essentially absent from both the granulocyte (Figure 1C) and B-cell (Figure 1D) compartments. These findings suggest that a subset of host CD8^+^ T cells, most likely resident VM cells present before irradiation, survived the conditioning regimen and persisted long term. Notably, VM cells were a small minority among CD8^+^ T cells derived from either young BM donors (Figure 1E) or aged BM donors (Figure 1F). In contrast, the majority of host-derived CD8^+^ T cells displayed a VM phenotype (Figure 1G). These findings suggest that VM cells generated from adult HSCs contribute minimally to the VM compartment observed in aged mice and further imply that a large fraction of aged VM cells originates early in life.

**Figure 1.**
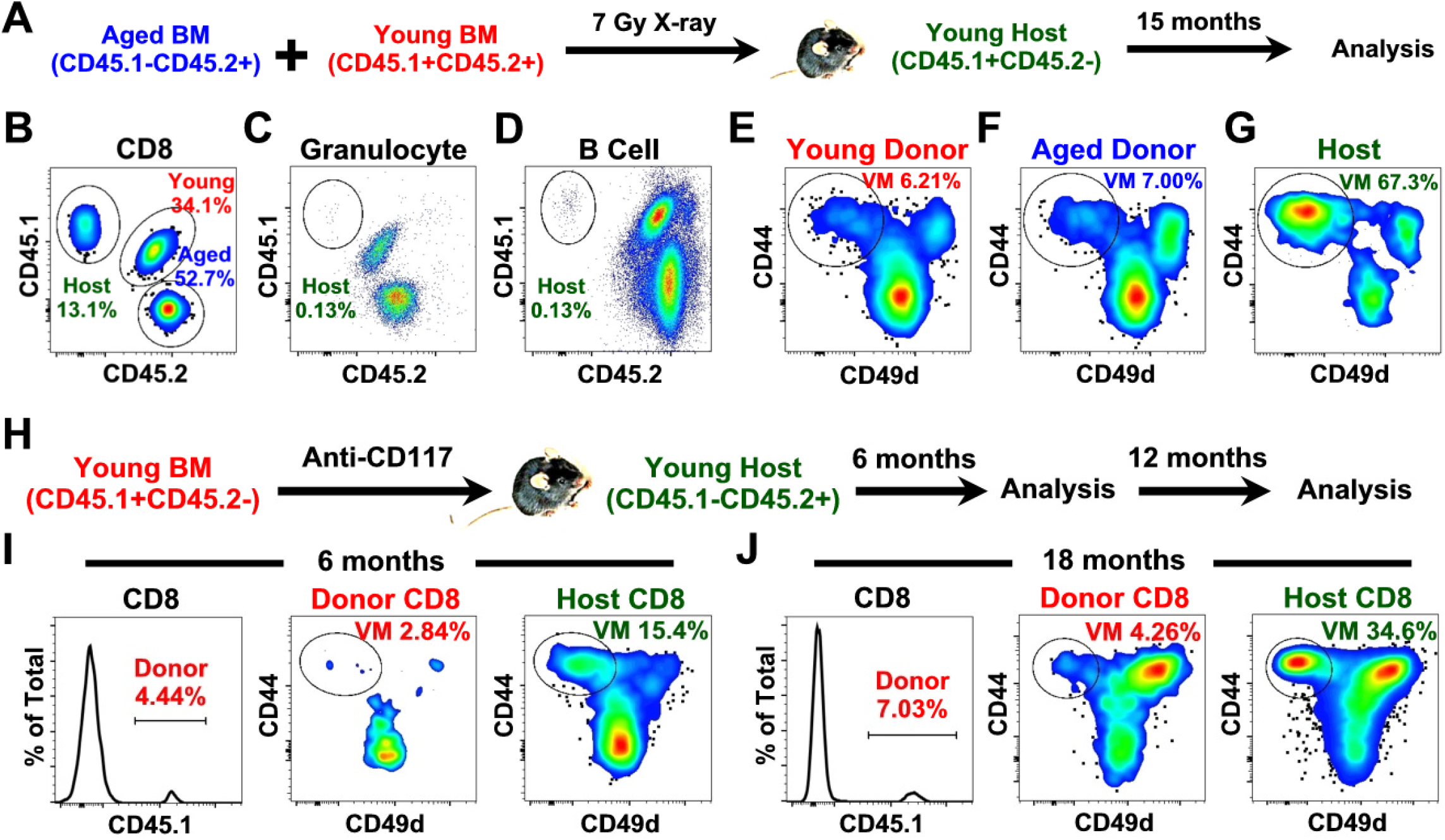
VM cells in aged mice arise predominantly early in life. (A-G) Congenically marked bone marrow (BM) from young (CD45.1+CD45.2+) and aged (CD45.1-CD45.2+) mice were transferred into x-ray irradiated young recipients (CD45.1+CD45.2-). Recipient mice were aged for 15 months before analysis. (A) Experimental design. (B-D) Young-BM-derived, aged-BM-derived and host cells were identified in CD8 T (B), granulocyte (C) and B cell (D) population based on their expression of the congenic markers. (E-G) VM cells were identified in young-BM-derived (E), aged-BM-derived (F) and host (G) CD8 T cell populations. Data are representative of 3 independent experiments. (H-J) Congenically marked (CD45.1-CD45.2+) young (2-month-old) mice were treated with anti-CD117 injection. Eight and 10 days later, congenically marked (CD45.1+CD45.2-) bone marrow (BM) cells were isolated from young (2-month-old) mice, depleted of T cells and transferred to the anti-CD117 treated mice. Six and 18 months later, peripheral blood samples were collected from the recipients and analyzed by flow cytometry. (H) Experimental design. At 6 months (I) and 18 months (J) post BM cell transfer, donor-BM-derived CD8 T cells were identified based on their expression of CD45.1, VM cells were identified in the donor-BM-derived and host CD8 T cells. Data are representative of 4 independent experiments.

To evaluate the contribution of adult thymopoiesis without the confounding effects of irradiation, congenically marked BM cells (CD45.1^+^) from young adult mice (2 months old) were transferred into age-matched recipients treated with anti-CD117, a conditioning strategy that permits HSC engraftment without irradiation [19] (Figure 1H). Peripheral blood samples were collected and analyzed by flow cytometry 6 and 18 months after transplantation. Donor-derived CD8^+^ T cells were identified by expression of CD45.1 (Figures 1I and 1J). The frequency of donor-derived cells remained stable between 6 and 18 months after transplantation, indicating durable engraftment of donor HSCs. Despite long-term reconstitution, donor-derived CD8^+^ T cells contained very few VM cells at either time point. In contrast, the frequency of VM cells within the host-derived CD8^+^ T-cell compartment increased markedly between 6 and 18 months after transplantation (Figures 1I and 1J), consistent with the known age-associated expansion of VM cells [15]. Together, these results indicate that the VM compartment in aged mice is derived predominantly from cells generated early in life, whereas adult thymopoiesis makes only a limited contribution.

### Resident VM cells are maintained through self-renewal with minimal input from naïve CD8^+^ T cells

Recent studies have shown that the naïve CD8^+^ T-cell compartment is largely depleted of VM precursors [3]. However, earlier reports demonstrated that naïve CD8^+^ T cells can give rise to VM cells under steady-state conditions [4]. To directly assess the contribution of naïve CD8^+^ T cells to VM-cell generation, congenically marked total CD8^+^ T cells (CD45.1^+^ CD45.2^+^) and purified naïve CD8^+^ T cells (CD44^lo^, CD45.1^+^ CD45.2^−^) were co-transferred into age-matched recipients (CD45.1^−^CD45.2^+^) (Figure 2A). Prior to transfer, donor cells were labeled with CFSE to monitor cell division.

**Figure 2.**
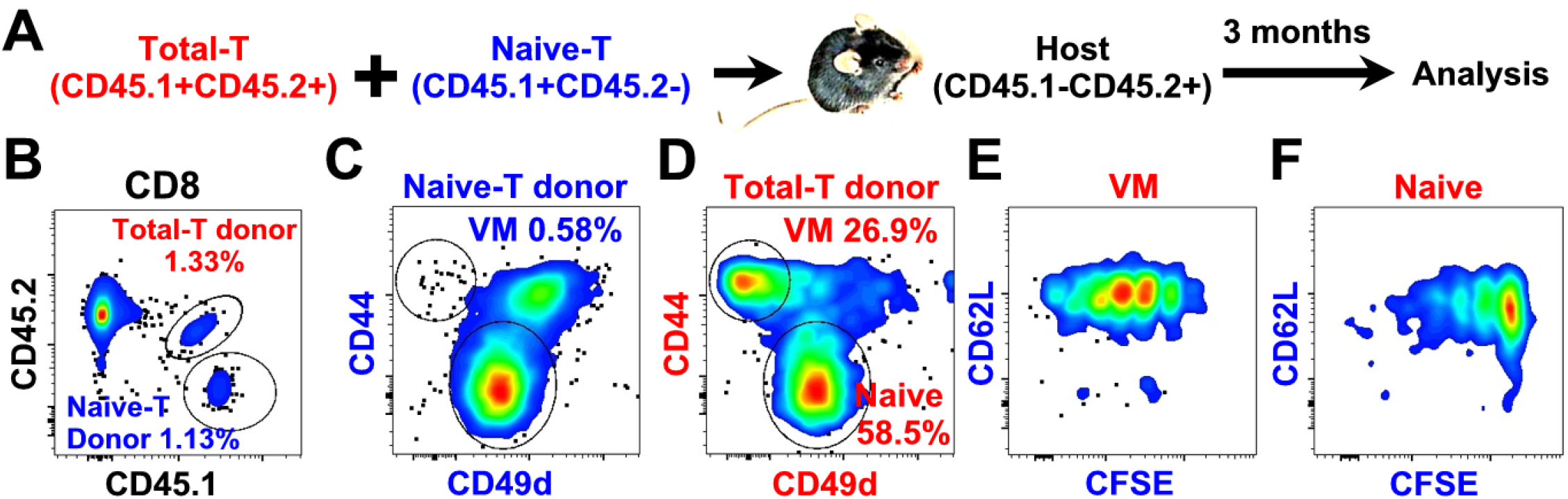
Resident VM cells are maintained through self-renewal with minimal input from naïve CD8^+^ T cells. Congenically marked total (CD45.1+CD45.2+) and naïve (CD44 low, CD45.1+CD45.2-) CD8 T cells were isolated from young (2-4 months) mice, mixed at 1:1 ratio, labeled with CFSE and transferred to age-matched hosts (CD45.1-CD45.2+). Three months later, donor cells were recovered and analyzed using flow cytometry. (A) Experimental design. (B) Total-T donor and naïve-T donor cells were identified based on their expression of the congenic markers (CD45.1 and CD45.2). (C) VM cells were identified in the naïve-T donor-derived cells. (D) VM and naïve cells were identified in total-T donor-derived cells. (E and F) CD62L expression levels and CFSE profiles of the VM (E) and naïve (F) cells of the total-T donor-derived cell population were determined. Data are representative of 3 independent experiments.

Three months after transfer, donor-derived cells were recovered and analyzed by flow cytometry. The two donor populations were readily distinguished by their congenic markers (Figure 2B). Strikingly, donor naïve CD8^+^ T-cell generated virtually no VM cells during the observation period (Figure 2C), consistent with the idea that the adult naïve CD8^+^ T-cell compartment contains very few VM precursors. In contrast, transferred total CD8^+^ T cells gave rise to a substantial population of VM cells (Figure 2D). Importantly, all donor-derived VM cells exhibited CFSE dilution (Figure 2E), indicating that they had undergone cell division during the 3-month period. Most VM cells divided approximately three to four times, although considerable variation was observed among individual cells, with some dividing only once and others undergoing as many as seven divisions. These findings suggest that peripheral VM cells are maintained through continuous self-renewal and reveal substantial heterogeneity in their proliferative behavior. In contrast, most donor-derived naïve CD8^+^ T cells remained undivided (Figure 2F). Both VM and naïve populations retained CD62L expression throughout the experiment (Figures 2E and 2F).

Together, these data indicate that adult naïve CD8^+^ T cells contribute minimally to VM-cell generation under steady-state conditions and support a model in which the peripheral VM-cell compartment is maintained primarily through self-renewal of pre-existing VM cells. In addition, the broad distribution of division histories suggests that VM cells differ substantially in their capacity for homeostatic self-renewal.

### The competitive fitness of the VM-cell compartment increases with age and is associated with enrichment of CD122hi cells

To directly compare the competitive fitness of young and aged VM cells in vivo, congenically marked CD8 +T cells from young (2–4 months; CD45.1^+^ CD45.2^−^) and aged (18–20 months; CD45.1^−^CD45.2^+^) donors were co-transferred into young recipients (CD45.1^+^ CD45.2^+^) (Figure 3A). Peripheral blood samples were collected 1 day, 6 months, and 12 months after transfer to determine the relative contribution of young and aged donor-derived VM cells. As expected, the frequency of VM cells within the CD8^+^ T-cell compartment increased with age (Figures 3B–3D). One day after transfer, aged donor-derived VM cells were only modestly more abundant than their young counterparts, with less than a twofold difference in frequency (Figure 3B). However, this disparity increased progressively over time. Whereas the number of aged donor-derived VM cells increased substantially, the number of young donor-derived VM cells declined (Figures 3B–3D). By 12 months after transfer, the ratio of aged to young donor-derived VM cells had reached approximately 30:1 (Figure 3D). These findings indicate that aged VM cells possess a marked competitive advantage over young VM cells and suggest that the overall fitness of the VM-cell compartment increases with age.

**Figure 3.**
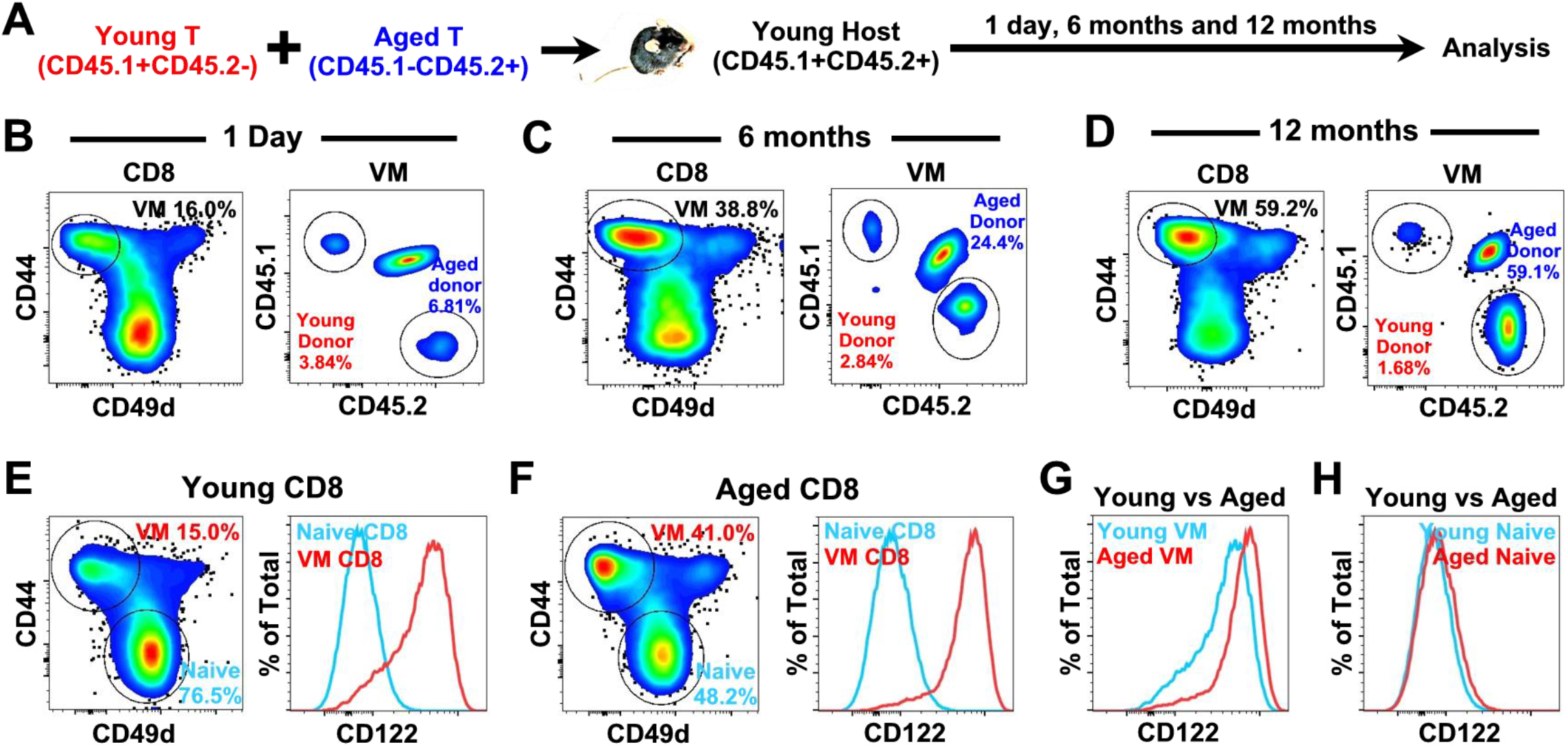
The competitive fitness of the VM-cell compartment increases with age and is associated with enrichment of CD122hi cells. Congenically marked T cells were isolated from young (2-4 months, CD45.1+CD45.2-) and aged (18-20 months, CD45.1-CD45.2+) mice and transferred to young (2-4 months) mice (CD45.1-CD45.2+). Peripheral blood samples were recovered and analyzed 1 day, 6 months and 12 months later using flow cytometry. (A) Experimental design. (B-D) Peripheral blood samples were analyzed at 1 day (B), 6 months (C) and 12 months (D) post cell transfer. The percentage of VM cells in the CD8 T cell population and the representation of young donor and aged donor-derived cells in the VM cell population were determined. Data are representative of 3 independent experiments. (E-G) CD8 T cells from young (4-month-old) and aged (20-month-old) were compared. Naïve and VM cells in CD8 T cell population in young (E) and aged (F) mice were identified and the levels of CD122 expression on naïve and VM CD8 T cells were compared. (G) The levels of CD122 expression on VM cells in young and aged mice were compared. Data are representative of 2 independent experiments.

To determine if the VM-cell compartment undergoes phenotypic changes with age, we focused on IL-15 responsiveness. VM-cell survival and homeostatic expansion depend on IL-15 [20]. Responsiveness to IL-15 is mediated by the β (CD122) and common γ (CD132) chains of the IL-15 receptor [21]. Whereas CD132 is constitutively expressed by T cells, CD122 expression varies substantially among T-cell subsets [22]. Consistent with previous studies [23], VM cells expressed higher levels of CD122 than naïve CD8^+^ T cells on average. However, considerable heterogeneity in CD122 expression was observed within the VM-cell compartment itself (Figure 3E), with substantial overlap between naïve and VM populations. This heterogeneity is consistent with the broad range of proliferative histories observed among VM cells (Figure 2E) and suggests that individual VM cells may differ in their capacity to respond to IL-15. If access to IL-15 is limiting under steady-state conditions, cells expressing higher levels of CD122 would be predicted to possess a competitive advantage and gradually become enriched within the VM-cell compartment. To test this possibility, CD122 expression was compared between VM cells from young and aged mice. Although substantial heterogeneity remained evident in aged animals, the distribution of CD122 expression was narrower than that observed in young mice (Figure 3F). Moreover, the VM-cell population underwent a clear age-associated shift toward higher CD122 expression (Figure 3G). These data demonstrate that the VM-cell compartment is not static. Its phenotypic composition shifts over time, including progressive enrichment of CD122^hi^ cells, accompanied by an increase in overall competitive fitness. Together, these findings indicate that the VM-cell compartment is a dynamic population that undergoes progressive phenotypic evolution throughout life as a consequence of continuous self-renewal and competition among resident cells.

### Young donor VM cells fail to expand efficiently in aged hosts

To determine whether age-associated changes in the host environment contribute to VM-cell expansion, congenically marked CD8^+^ T cells from young mice (2 months old; CD45.1^+^ CD45.2^−^) were transferred into either young (2 months old) or aged (20 months old) recipients (CD45.1^−^CD45.2^+^) (Figure 4A). Donor cells were labeled with CFSE prior to transfer to monitor cell division. Peripheral blood samples were collected from donors and recipients before transfer to establish baseline frequencies of VM cells. As expected, the frequency of VM cells was substantially lower in young donors (Figure 4B) and young recipients (Figure 4C) than in aged recipients (Figure 4D). Three months after transfer, donor-derived cells were recovered and analyzed by flow cytometry. Donor cells were identified based on expression of the congenic marker CD45.1 (Figures 4E and 4F).

**Figure 4.**
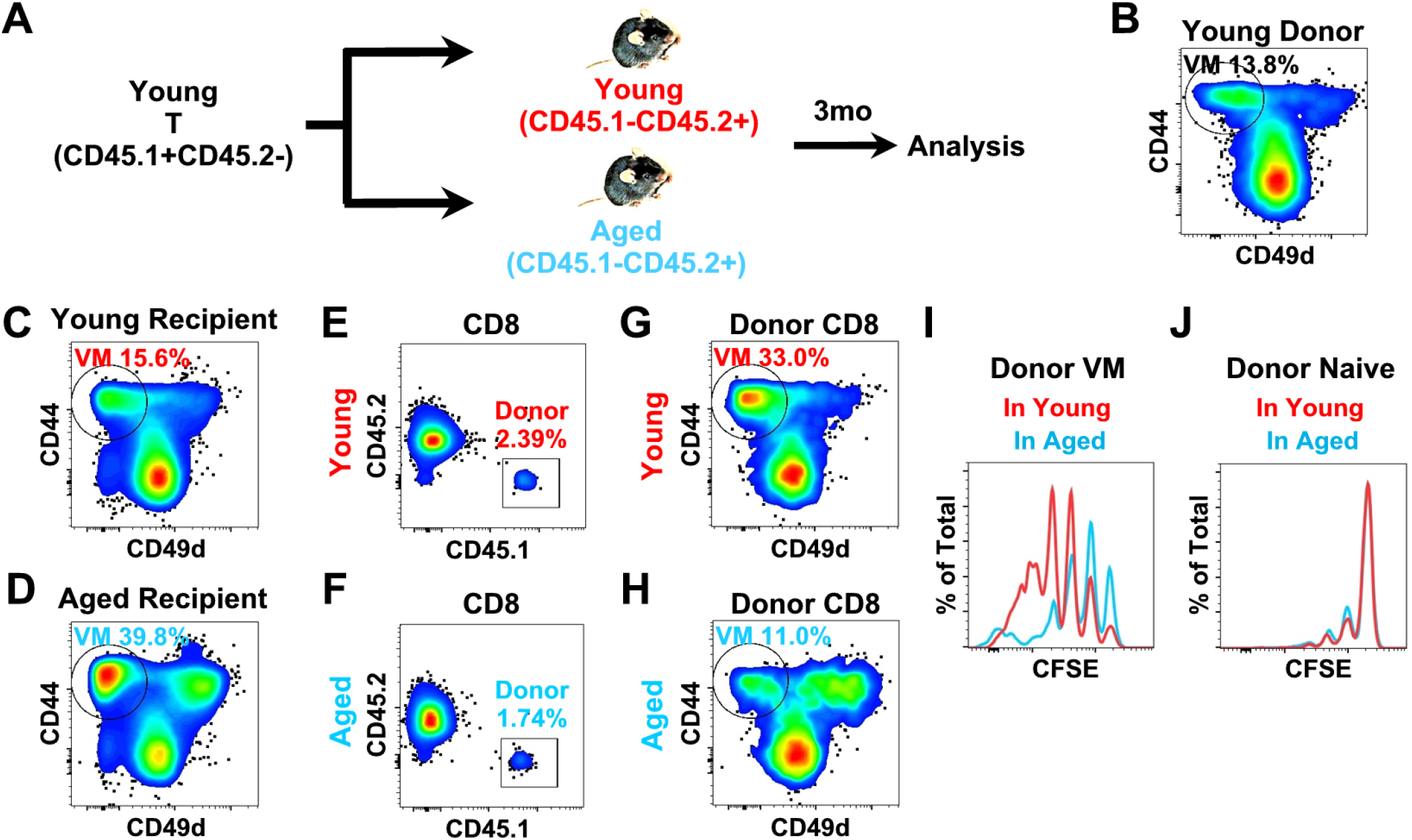
Young donor VM cells fail to expand efficiently in aged hosts. Congenically marked (CD45.1+CD45.2^−^) T cells were isolated from young (2-4 months) mice, labeled with CFSE and transferred to young (2-4 months) or aged (18-20 months) mice (CD45.1-CD45.2+). Three months later, donor cells were recovered and analyzed using flow cytometry. (A) Experimental design. (B-D) Peripheral blood samples from the young donors (B), the young recipients (C) and the aged recipients (D) were analyzed for the percentage of VM cells in the CD8 T cell population before cell transfer. (E and F) Donor cells in CD8 T cell population in young (E) and aged (F) recipients were identified based on their expression of the congenic markers (CD45.1). (G and H) VM cells were identified in the donor CD8 T cell population in young (G) and aged (H) recipients. (I and J) Donor VM (I) and naïve (J) cells recovered from the young and aged recipients were compared for their CFSE profiles. Data are representative of 3 independent experiments.

In young recipients, the frequency of VM cells within the donor-derived CD8^+^ T-cell population increased markedly during the 3-month observation period (Figure 4G compared to 4C). In contrast, donor-derived VM cells failed to expand in aged recipients, and their frequency remained unchanged or decreased slightly over the same period (Figure 4H compared to 4D). Consistent with these findings, donor-derived VM cells underwent substantially more cell division in young recipients than in aged recipients, as assessed by CFSE dilution (Figure 4I). In contrast, donor-derived naïve CD8^+^ T cells displayed similar division profiles in young and aged recipients (Figure 4J). Together, these findings indicate that age-associated expansion of the VM-cell compartment is not driven by changes in the host environment. Rather, aged host environments appear less permissive for VM-cell expansion and self-renewal than young host environments. One possible explanation is that resident VM cells in aged mice occupy limiting homeostatic niches required for VM-cell maintenance and expansion, thereby restricting the accumulation of newly generated VM cells.

### Reduced TGFβ signaling enhances the competitive fitness of VM cells

Expression of CD122 on memory-phenotype CD8^+^ T cells is positively regulated by IL-15 and negatively regulated by TGFβ [24]. To further investigate the role of CD122 in regulating VM-cell fitness, VM cells from transgenic mice expressing a dominant-negative TGFβ receptor II (TGFβRDN) were examined. Previous studies reported accelerated expansion of memory-phenotype (CD44^high) CD8^+^ T cells in TGFβRDN mice [25, 26], although it remained unclear whether the expanded population consisted primarily of VM cells. To address this question, CD8^+^ T cells from young (2-month-old) TGFβRDN mice were compared with those from age-matched wild-type (WT) controls. As shown in Figure 5A, the frequency of VM cells was more than twofold higher in TGFβRDN mice than in WT controls, indicating that reduced TGFβ signaling promotes expansion of the VM-cell compartment.

**Figure 5.**
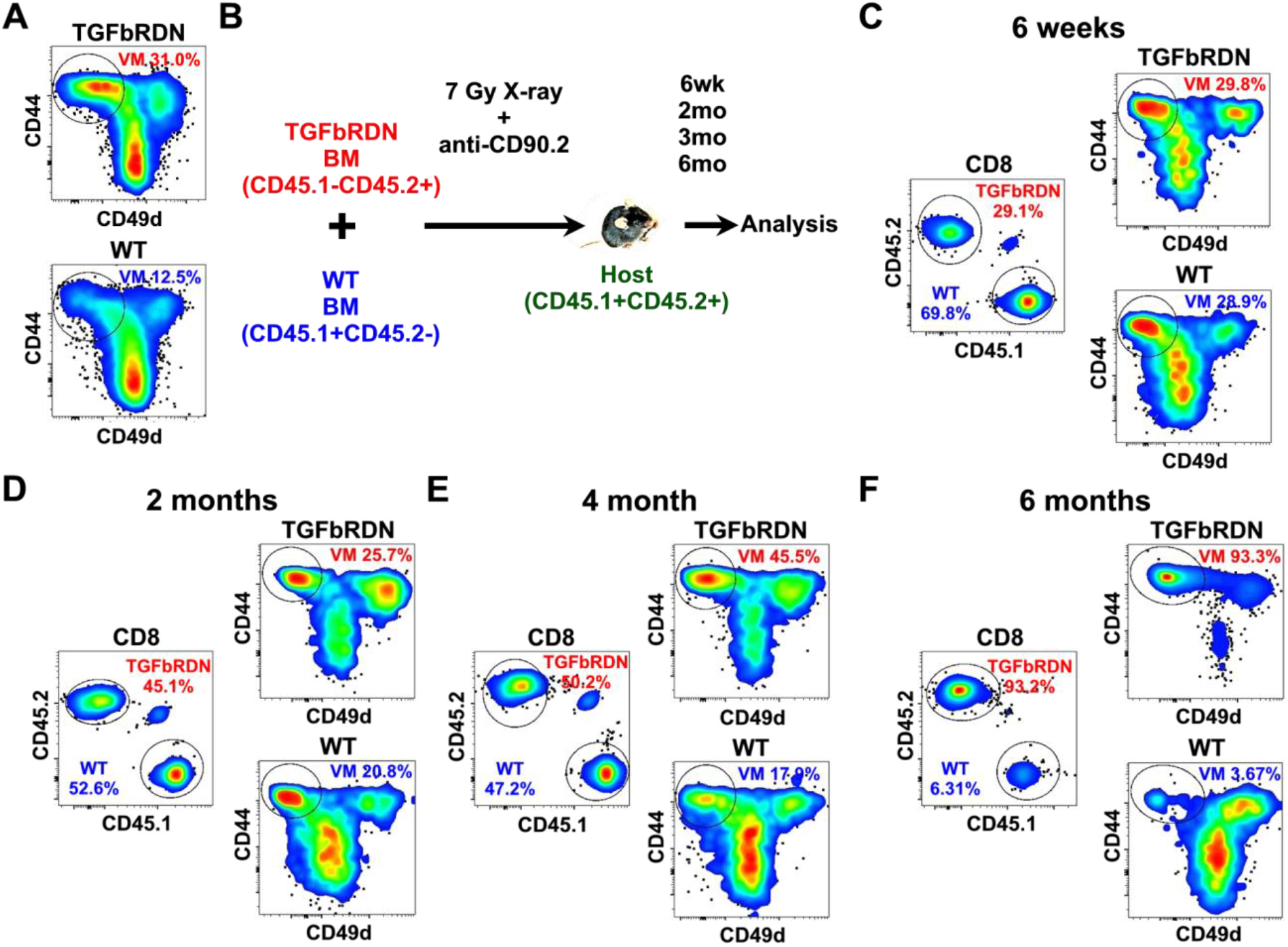
Reduced TGFβ signaling enhances the competitive fitness of VM cells. (A) CD8 T cells from young (2-month-old) TGFbRDN transgenic and age-matched WT mice were analyzed by flow cytometry. VM cells were identified in the CD8 T cell population. (B-F) Congenically marked bone marrow cells (BM) from TGFbRDN transgenic (CD45.1-CD45.2+) and WT (CD45.1+CD45.2-) mice were mixed at 1 to 2 ratio and transferred into x-ray irradiated and T cell depleted young recipients (CD45.1+CD45.2+). Peripheral blood samples were collected and analyzed at 6 weeks, 2 months, 4 months and 6 months post bone marrow reconstitution. (B) Experimental design. (C-F) At 6 weeks (C), 2 months (D), 4 months (E) and 6 months (F) post bone marrow reconstitution, TGFbRDN transgenic BM-derived cells and WT BM-derived cells were identified in CD8 T cell population based on the expression of their congenic markers, and VM cells were identified in the TGFbRDN transgenic BM-derived CD8 and WT BM-derived CD8 T cell populations. Data are representative of 2 independent experiments.

To compare TGFβRDN and WT cells within the same environment, congenically marked bone marrow cells from TGFβRDN mice (CD45.1^−^CD45.2^+^) and WT mice (CD45.1^+^ CD45.2^−^) were co-transferred into irradiated, T-cell-depleted WT recipients (CD45.1^+^ CD45.2^+^) (Figure 5B). Preliminary experiments showed that equal mixtures of WT and TGFβRDN bone marrow generated overwhelming dominance of TGFβRDN-derived VM cells. Therefore, subsequent experiments were performed using a 2:1 excess of WT bone marrow cells to facilitate analysis of both populations. Peripheral blood samples were collected 6 weeks, 2 months, 4 months, and 6 months after reconstitution. At 6 weeks, the ratio of TGFβRDN-derived to WT-derived CD8^+^ T cells closely reflected the input bone marrow ratio (Figure 5C). In addition, the frequencies of VM cells within the TGFβRDN-derived and WT-derived CD8^+^ T-cell populations were comparable. These findings suggest that reduced TGFβ signaling has little effect on the initial generation of CD8^+^ T cells or VM cells.

In contrast, striking differences emerged over time. The proportion of TGFβRDN-derived CD8^+^ T cells increased progressively relative to WT-derived cells (Figures 5D–5F). By 6 months after reconstitution, TGFβRDN-derived cells accounted for more than 90% of the total CD8^+^ T-cell compartment (Figure 5F). Consistent with previous reports [25, 26], the chimeric mice developed lymphoproliferative disease. The VM-cell compartment exhibited an even more pronounced bias. The frequency of VM cells increased progressively within the TGFβRDN-derived CD8^+^ T-cell population, whereas it declined within the WT-derived population. By 6 months after reconstitution, more than 90% of TGFβRDN-derived CD8^+^ T cells displayed a VM phenotype, while WT-derived VM cells were nearly absent (Figure 5F).

Together, these findings demonstrate that reduced TGFβ signaling confers a marked long-term competitive advantage within the VM-cell compartment. Given that reduced TGFβ signaling only transiently and moderately increases the levels of CD122 [24], these results are consistent with a model in which relatively small variations in competitive fitness can ultimately lead to progressive enrichment and expansion of the most competitive VM cells.

### Long-term self-renewal promotes the emergence of dominant clonal VM-cell populations

Previous studies showed that memory-phenotype CD8^+^ T cells undergo progressive expansion and after a few months, give rise to dominant clonal populations in both TGFβRDN [25, 26] and IL-15 transgenic [27] mice. Because VM cells constitute the major memory-phenotype CD8^+^ T-cell population under steady-state conditions, these observations suggest that VM cells may be the source of these clonal populations. The data presented in Figure 5 further support this possibility by demonstrating that enhanced competitive fitness within the VM-cell compartment results in progressive expansion and eventual lymphoproliferative disease. To determine whether prolonged self-renewal of VM cells can promote the emergence of dominant clonal populations, a serial transfer strategy was employed. As shown in Figure 6A, congenically marked T cells from aged mice were transferred into young recipients. Twelve months later, donor-derived cells were recovered and transferred into a second cohort of young recipients. The process was repeated once more to generate tertiary recipients. Strikingly, all tertiary recipients, approximately 60% of secondary recipients, and 10% of primary recipients developed splenomegaly and lymphadenopathy before completion of the 12-month observation period. In each case, disease was associated with massive expansion of donor-derived VM cells. To assess clonality, donor-derived VM cells from affected animals were analyzed using a panel of Vβ-specific antibodies. As shown in Figure 6B, the expanded populations were highly clonal. In the representative example shown, essentially all donor-derived VM cells expressed Vβ8.1/8.2, indicating marked enrichment of a single clone.

**Figure 6.**
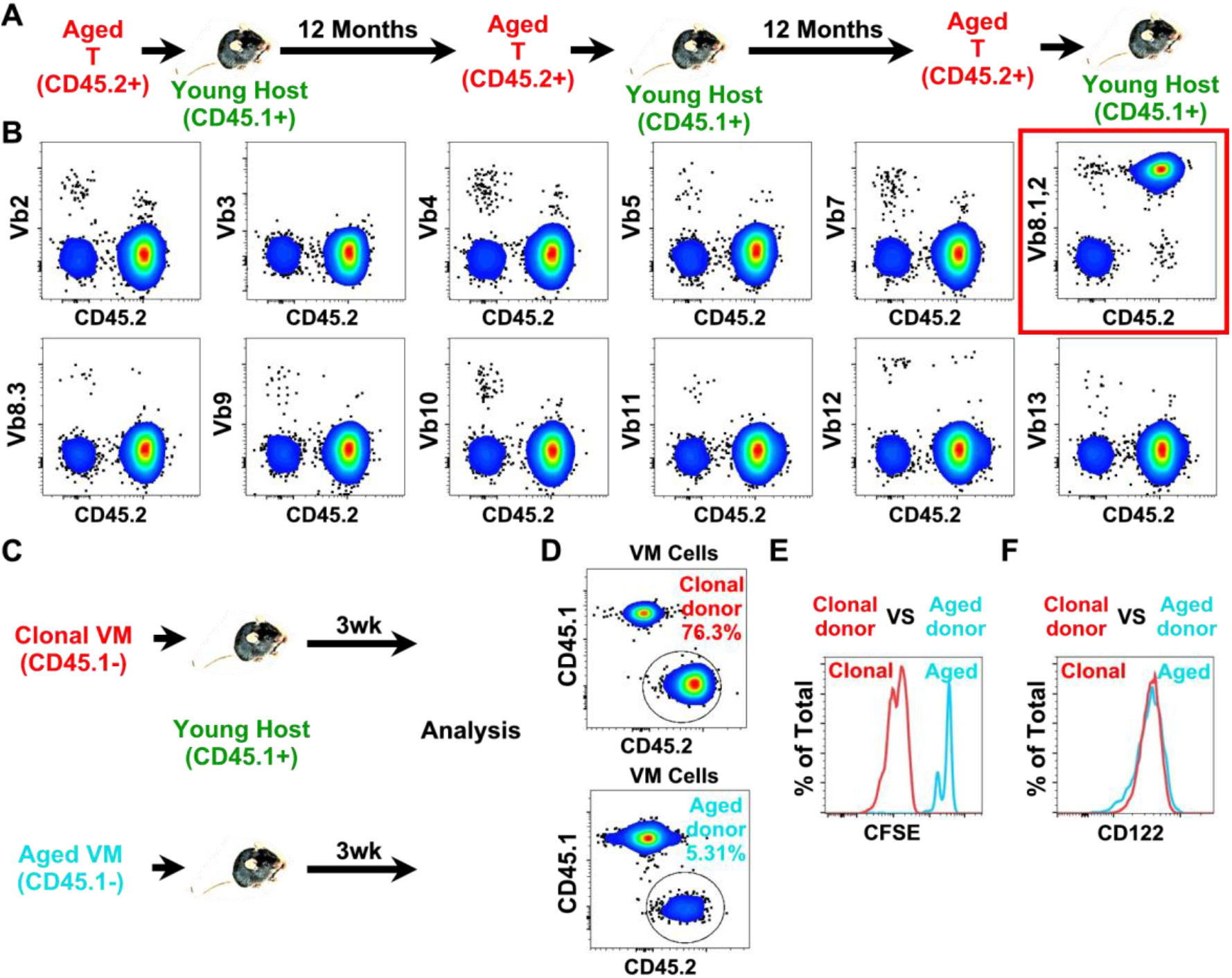
Long-term self-renewal promotes the emergence of dominant clonal VM-cell populations. (A-B) Congenically marked (CD45.1-CD45.2+) T cells were isolated from aged (20-month-old) mice and transferred to young (2-month-old, CD45.1+CD45.2-) hosts. Twelve months later, donor cells were recovered and transferred to a new set of young (2-month-old, CD45.1+CD45.2-) hosts. Twelve months later, donor cells were recovered and transferred again to a new set of young (2-month-old, CD45.1+CD45.2-) hosts. (A) Experimental design. (B) CD8 T cells from VM recipients that developed splenomegaly were analyzed by flow cytometry for their expression of the congenic marker CD45.2 and the indicated Vb chain of the T cell receptor. Data are representative of 3 independent experiments. (C-F) T cells from VM cell recipients that developed splenomegaly were labeled with CFSE and transferred to congenically marked (CD45.1+CD45.2-) young hosts. Three weeks later, VM cells were recovered from the recipients and analyzed by flow cytometry. In control experiments, T cells from aged mice were labeled with CFSE and transferred to congenically marked (CD45.1+CD45.2-) young hosts. Three weeks later, VM cells were recovered from the recipients and analyzed by flow cytometry. (C) Experimental design. (D) Donor cells and host cells in the VM cell population were identified based on their expression of the congenic marker. (E) The CFSE profiles of the clonal VM donors and the aged VM donors were compared. (F) The CD122 expression levels of the clonal VM donors and the aged VM donors were compared. Data are representative of 2 independent experiments.

To determine whether these clonal VM populations exhibited altered proliferative behavior, clonal VM cells were isolated, labeled with CFSE, and transferred into congenically marked young recipients (CD45.1^+^). VM cells isolated from normal aged mice were analyzed in parallel as controls (Figure 6C). Three weeks after transfer, donor-derived cells were recovered and analyzed by flow cytometry. Recipients of clonal VM cells contained more donor-derived VM cells than host VM cells after only three weeks (Figure 6D). In contrast, donor-derived VM cells from normal aged mice accounted for only a small fraction of the total VM-cell compartment. Consistent with these findings, clonal VM cells underwent extensive proliferation, dividing more than four times during the 3-week period (Figure 6E). In comparison, only a small subset of control VM cells divided once, whereas the majority remained undivided. Despite their dramatically different proliferative behavior, clonal and control VM cells expressed similar levels of CD122 (Figure 6F), indicating that the enhanced growth of the clonal populations could not be explained solely by increased IL-15 responsiveness.

Together, these findings demonstrate that prolonged self-renewal of VM cells is associated with the emergence of highly competitive clonal populations that exhibit accelerated proliferation and progressive dominance in vivo. Notably, unlike the fitness advantages described in Figures 3–5, the enhanced growth of these clonal populations was not accompanied by increased CD122 expression, suggesting that they appear to be less constrained by the homeostatic mechanisms that regulate typical VM-cell populations.

### Removal of resident VM cells resets the aged VM-cell compartment

To directly assess the role of resident VM cells in maintaining age-associated expansion of the VM-cell compartment, depleting antibodies were used to eliminate VM cells in young (6-month-old) and aged (18-month-old) mice. Several antibodies were evaluated, including anti-CXCR3 (clone CXCR3-137), anti-Ly6C (clone HK1.4), anti-CD122 (clone TM-b1), and anti-Gr1 (clone RB6-8C5). Complete depletion of VM cells was achieved only with anti-CD122 and anti-Gr1. Because substantially lower doses of anti-Gr1 were required to achieve depletion, anti-Gr1 was used in subsequent experiments (Figure 7A). In young mice, anti-Gr1 treatment effectively depleted VM cells, leaving very few detectable cells 10 days after treatment (Figure 7B). As expected, VM-cell frequencies remained unchanged in control IgG-treated animals. Following depletion, the VM-cell compartment recovered rapidly. Two months after treatment, VM-cell frequencies had nearly returned to those observed in control animals (Figure 7C). Both depleted and control groups subsequently exhibited progressive age-associated expansion of the VM-cell compartment, reaching similar frequencies 5 (Figure 7D) and 12 (Figure 7E) months after treatment. These findings indicate that, in the absence of pre-existing resident VM cells, newly generated VM cells arising in adulthood retain the capacity to expand and establish a normal age-associated VM-cell compartment.

**Figure 7.**
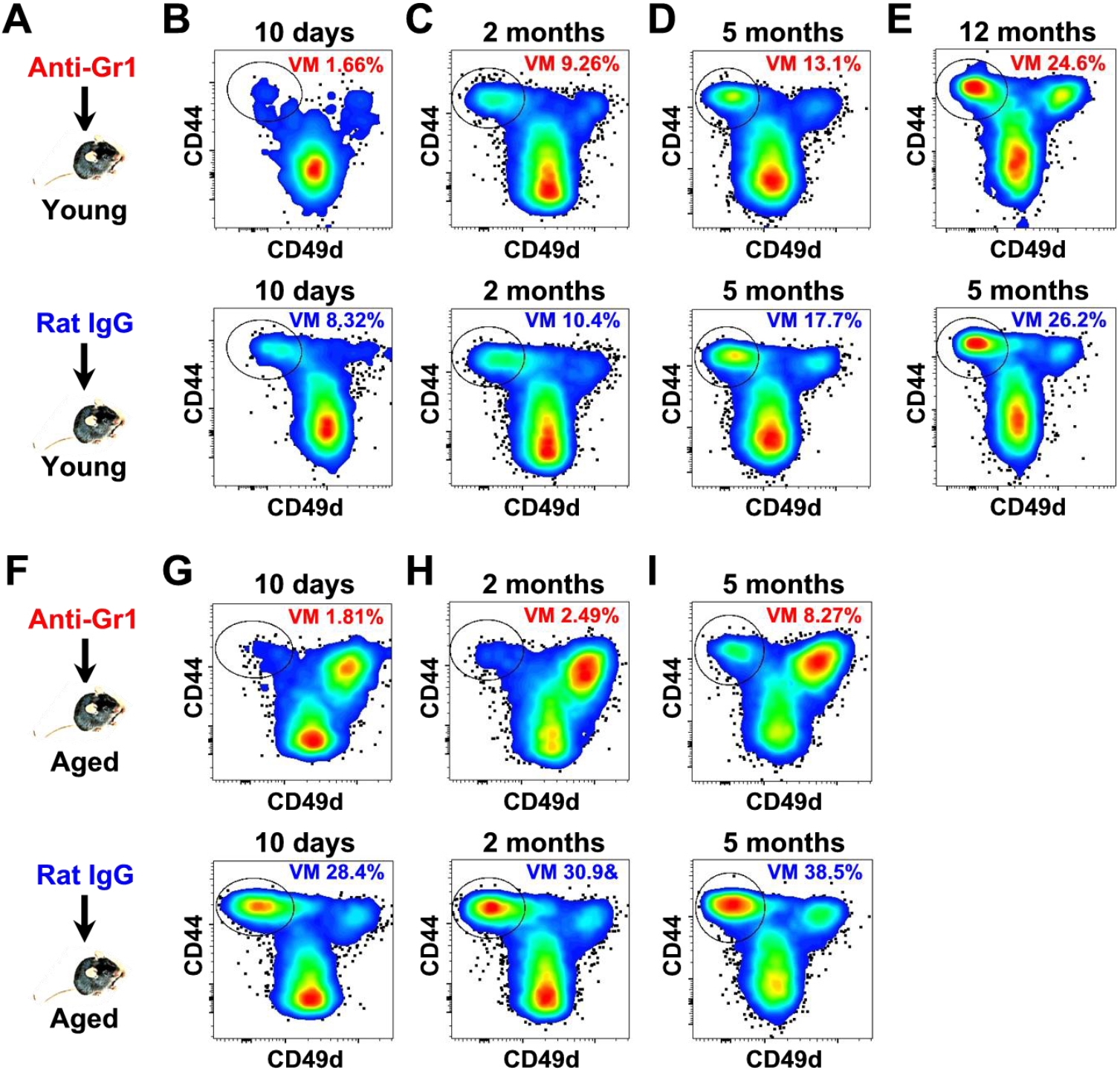
Removal of resident VM cells resets the aged VM-cell compartment. (A–E) Young (6-month-old) mice were treated with anti-Gr1 antibody (clone RB6-8C5) to deplete VM cells or with rat IgG as a control. (A) Experimental design. Peripheral blood samples were collected and analyzed by flow cytometry 10 days (B), 2 months (C), 5 months (D), and 12 months (E) after treatment. VM cells were identified within the CD8^+^ T-cell population, and the percentages of VM cells in anti-Gr1-treated and rat IgG-treated mice were compared. (F–I) Aged (18-month-old) mice were treated and analyzed as described in (A–E). (F) Experimental design. Peripheral blood samples were collected and analyzed by flow cytometry 10 days (G), 2 months (H), and 5 months (I) after treatment. VM cells were identified within the CD8^+^ T-cell population, and the percentages of VM cells in anti-Gr1-treated and rat IgG-treated mice were compared. Data are representative of two independent experiments.

A similar depletion strategy was applied to aged mice (Figure 7F). As in young animals, anti-Gr1 treatment efficiently eliminated VM cells, whereas control IgG treatment had little effect on the expanded VM-cell compartment characteristic of aged mice (Figure 7G). Strikingly, depletion of resident VM cells largely reversed the age-associated expansion phenotype. Recovery of the VM-cell compartment occurred substantially more slowly in aged mice than in young mice. Two months after depletion, only a small VM-cell population was detectable in aged animals (Figure 7H), and approximately 5 months were required for VM-cell frequencies to reach levels observed after only 2 months in young mice (Figure 7I compared with Figure 7C). Despite the slower recovery kinetics, the VM-cell compartment in aged mice ultimately re-established at frequencies typically observed in young adult animals rather than those characteristic of aged mice. These findings indicate that newly generated VM cells remain capable of expanding in aged animals and that age-associated VM-cell expansion is not maintained independently of the resident VM-cell population. Instead, the expanded VM-cell compartment characteristic of aging appears to depend on the continued persistence of long-lived resident VM cells.

## Discussion

VM cells represent a unique population of autoreactive CD8^+^ T cells that acquire a memory phenotype in the absence of foreign antigen exposure. Although age-associated expansion of VM cells has been recognized for many years, the mechanisms responsible for this phenomenon have remained poorly understood. In the present study, we show that the VM-cell compartment in aged mice is derived predominantly from cells generated early in life and maintained through continuous self-renewal. Over time, the VM-cell compartment becomes enriched for cells with greater competitive fitness and undergoes progressive phenotypic remodeling, including increased expression of CD122. These findings support a model in which long-term competition for limiting homeostatic resources progressively reshapes the VM-cell compartment during aging.

A central finding of this study is that most VM cells found in aged mice originate early in life. In both irradiation-based and anti-CD117-mediated bone marrow reconstitution models, adult-derived hematopoietic stem cells contributed very little to the aged VM-cell compartment despite long-term engraftment. These observations indicate that the expanded VM-cell population characteristic of aged mice is not generated continuously throughout life but instead derives largely from cells established during early development. This conclusion is consistent with previous studies showing that CD8^+^ T cells generated during the neonatal period exhibit a substantially greater propensity to adopt the VM-cell fate than those generated during adulthood [17]. The persistence of early-life VM cells raised the question of how these cells are maintained throughout life. Our adoptive transfer experiments demonstrate that the VM-cell compartment is maintained primarily through self-renewal rather than continuous recruitment from the adult naïve CD8^+^ T-cell pool. Moreover, VM cells exhibited substantial heterogeneity in their proliferative histories, suggesting that individual VM cells differ considerably in their capacity for homeostatic self-renewal. This heterogeneity ultimately provided an important clue to the mechanism underlying age-associated VM-cell expansion.

Several lines of evidence suggest that differences in responsiveness to homeostatic signals play an important role in shaping the VM-cell compartment. First, VM cells displayed substantial heterogeneity in CD122 expression, indicating phenotypic diversity within the population. Second, the VM-cell compartment became progressively enriched for CD122^hi^ cells with age. Third, aged VM cells exhibited a marked competitive advantage over young VM cells in long-term adoptive transfer experiments. Finally, experimental reduction of TGFβ signaling dramatically enhanced the competitive fitness of VM cells and resulted in progressive dominance of the VM-cell compartment. Together, these observations indicate that VM cells undergo phenotypic evolution during aging and that differences in responsiveness to homeostatic cues can profoundly influence long-term competitive fitness. Although CD122 expression provides one plausible mechanism underlying these differences, the present study does not establish that IL-15 is the sole determinant of VM-cell fitness. In addition to IL-15, other factors may contribute to the competitive interactions that shape the VM-cell compartment, including self-antigen recognition, IL-7 signaling, TGFβ signaling, or other homeostatic cues. Thus, we favor a model in which lifelong self-renewal allows gradual selection of VM cells with superior fitness, while the precise molecular determinants of that fitness remain to be fully defined.

It was a long-held belief that the lack of thymic production in aged mice leads to increased homeostatic proliferation of naïve T cells, which in turn convert naïve T cells into VM cells [28, 29]. This notion is supported by the fact that similar to age-related lymphopenia, neonatal lymphopenia elevates the frequency of VM cells [2, 4]. However, recent data showed that naïve and VM cells belong to different lineages [3]. Our data show that naïve CD8 T cells rarely give rise to VM cell under steady-state conditions. The observation that donor-derived VM cells expanded poorly in aged recipients initially suggested that age-associated changes in the host environment might limit VM-cell expansion. However, depletion of resident VM cells revealed a different explanation. Following removal of the resident VM-cell compartment, newly generated VM cells were capable of repopulating both young and aged animals. Although recovery occurred more slowly in aged mice, the aged VM-cell compartment was effectively reset and re-established at levels characteristic of young adults rather than aged animals. These findings indicate that newly generated VM cells remain capable of expansion in aged mice and suggest that the failure of these cells to accumulate under normal conditions results primarily from competition with long-lived resident VM cells. Thus, age-associated VM-cell expansion appears to be actively maintained by the resident population rather than by irreversible changes in the aging environment.

An unexpected finding was the emergence of dominant clonal VM-cell populations following prolonged serial propagation. The incidence of clonal expansion increased progressively with successive rounds of transfer, suggesting that extensive proliferative history promotes the development of highly competitive clones. These clonal populations exhibited accelerated proliferation and rapidly dominated recipient mice following transfer. Notably, their enhanced growth was not associated with increased CD122 expression, distinguishing them from the competitive advantages observed in non-transformed VM cells. This observation suggests that dominant clonal populations may eventually escape the homeostatic mechanisms that normally regulate VM-cell fitness. Although the molecular basis of this phenomenon remains unknown, the findings establish a direct link between long-term self-renewal and the emergence of highly competitive clonal populations within the VM-cell compartment.

The findings reported here may have implications beyond VM-cell biology. Aging is frequently accompanied by the expansion of memory-phenotype T cells and by increasing oligoclonality within the T-cell compartment. Similar phenomena have been observed in both mice and humans, yet the mechanisms responsible remain incompletely understood. More broadly, similar to VM cells, tissue-specific stem cells are maintained by self-renewal throughout life. Like VM cells, hematopoietic stem cells undergo population expansion and cellular senescence with aging [30, 31]. However, the underlying mechanism is poorly understood. The present study suggests that long-lived self-renewing cell populations may undergo progressive competitive selection throughout life, resulting in gradual enrichment of highly fit clones. Under some circumstances, this process may ultimately permit the emergence of dominant clonal populations with altered growth properties. Whether similar processes contribute to age-associated clonal expansions observed in other immune cell or tissue specific stem cell populations remains an important question for future investigation.

In summary, our findings identify a previously unrecognized mechanism underlying age-associated VM-cell expansion. Rather than reflecting continuous production of new VM cells or age-dependent changes in the host environment, the aged VM-cell compartment appears to result from lifelong self-renewal and competition among long-lived resident VM cells. This process progressively enriches cells with greater competitive fitness and, over extended periods, is associated with the emergence of dominant clonal populations. These observations provide a framework for understanding how aging reshapes autoreactive T-cell populations and reveal how long-term homeostatic competition can profoundly influence the composition of the immune system over time.

## Materials and Methods

### Mice

C57BL/6 mice 2–24 months of age were obtained from the National Institute on Aging contract colony at Harlan Laboratories (Indianapolis, IN) or from The Jackson Laboratory (Bar Harbor, ME). Congenic C57BL/6 mice, C57BL/6J (CD45.1^−^CD45.2^+^ ; Stock No. 000664) and B6.SJL-Ptprca Pepcb/BoyJ (CD45.1^+^ CD45.2^−^; Stock No. 002014), were purchased from The Jackson Laboratory. C57BL/6 F1 congenic mice (CD45.1^+^ CD45.2^+^) were generated by crossing male B6.SJL-Ptprca Pepcb/BoyJ (CD45.1^+^ CD45.2^−^) mice with female C57BL/6J (CD45.1^−^CD45.2^+^) mice. Mice were identified by ear tags and maintained under specific pathogen-free conditions with food and water provided ad libitum. All animal studies were approved by the University of Michigan Committee on the Use and Care of Animals (UCUCA).

### T cell adoptive transfer

Adoptive transfer experiments were performed as previously described [32]. Briefly, single-cell suspensions were prepared from freshly isolated spleens and lymph nodes by mechanical dissociation. Unwanted cell populations were depleted in vitro using biotinylated antibodies and anti-biotin negative-selection kits (Miltenyi Biotec, Auburn, CA). Cells were assessed for viability and transferred intravenously in 0.5 ml PBS. For long-term tracking experiments, at least 5 × 10^6^ viable cells were transferred.

### Generation of mixed bone marrow chimeras

To generate mixed bone marrow chimeras, congenically marked mice were irradiated with a single dose of 7 Gy and used as recipients. Lineage-depleted (CD4-, CD8-, CD5-, CD19-, B220-, NK1.1-, and CD11b-) bone marrow cells from congenically marked donors were mixed at a predetermined ratio, assessed for viability and transferred intravenously within 2 h after irradiation. In some cases, recipients were also injected intraperitoneally with 0.5 mg anti-CD90.2 (clone 30H12) antibody to deplete resident T cells.

Bone marrow chimeras were also generated without irradiation as previously described [33]. Briefly, recipient mice were injected i.p. with 0.5 mg anti-CD117 (ACK2) (eBioscience, San Diego, CA) in 0.5 ml PBS. Lineage-depleted (CD4-, CD8-, CD5-, CD19-, B220-, NK1.1-, and CD11b-) bone marrow cells were checked for viability and transferred i.v. in 0.3 ml PBS 8 and 10 days after anti-CD117 treatment.

### Flow cytometry

Flow cytometric analyses were performed as previously described [19]. Briefly, single-cell suspensions were prepared from peripheral blood, spleens, and lymph nodes. Spleen and lymph node suspensions were generated by mechanical dissociation. Peripheral blood samples were treated with red blood cell lysis buffer and washed prior to antibody staining.

Approximately 1 × 10^6^ cells were stained with fluorochrome-conjugated antibodies in 5% BSA-PBS buffer. Following staining, cells were washed with 1% BSA-PBS buffer and fixed/permeabilized using the Foxp3 Transcription Factor Fixation/Permeabilization Concentrate and Diluent kit (Thermo Fisher Scientific). Fixed and permeabilized cells were then stained with fluorochrome-conjugated antibodies in 5% BSA-PBS buffer. After washing, cells were resuspended in 300 μl of 1% BSA-PBS buffer and acquired on a BD LSR II flow cytometer. Data were analyzed using FlowJo software (Tree Star).

Antibodies used for flow cytometry included anti-CD19 (6D5), anti-CD62L (MEL-14), anti-CD44 (IM7), anti-CXCR3 (CXCR3-173), anti-TCRβ (H57-597), anti-CD49d (9C10), anti-CD122 (TM-b1), anti-CD5 (53-7.3), anti-CD6 (J90-462), anti-CD69 (H1.2F3), anti-CD127 (A7R34), anti-H-2K^b^ (28-8-6), anti-Ki-67 (SolA15), anti-CD4 (GK1.5), anti-CD11b/Mac-1 (M1/70), anti-CD45 (30-F11), anti-CD45.1 (A20), anti-CD45.2 (104) and anti-CD8b (H35-17.2). Antibodies were purchased from BioLegend (San Diego, CA), Serotec (Raleigh, NC), eBioscience (San Diego, CA), or Cell Signaling Technology and included eFluor 450-, APC-, APC-Cy7-, AF700-, PerCP-Cy5.5-, FITC-, PE-, PE-Cy7-, and PE-Cy5-conjugated antibodies.

## Notes

### Competing Interest Statement

The authors have declared no competing interest.

